# Treatment with valproate downregulates the *Agtr2* mRNA in rat lungs

**DOI:** 10.1101/2020.05.21.108399

**Authors:** Kadri Seppa, Anton Terasmaa, Toomas Jagomäe, Jürgen Innos, Eero Vasar, Mario Plaas

## Abstract

GLP1 receptor agonist liraglutide has been shown to upregulate ACE2 expressions in several animal studies and thereby mediate strong positive stress response^1^. On the other hand, two in silico studies suggest that valproate downregulates ACE2 and AGTR2 gene expressions^2,3^. In this study, we have evaluated how these two widely used drugs, liraglutide and valproate, change the expression pattern of RAS system genes in the rat lungs. Our results indicate that eight-day treatment with valproate significantly downregulates the gene expression of *Agtr2, Mas1* and *Agrt1b* in the rat lungs. These effects are reversed by co-administration of liraglutide.

## Introduction

The renin-angiotensin system (RAS) plays an important role in the control of body functions, including sodium and water content and blood pressure. Two main peptide hormones of the RAS system are the vasoconstrictor angiotensin II (Ang II) and the vasodilator angiotensin 1– 7 (Ang-(1–7)), which have several receptors: Ang II type 1 and type 2 receptors (AGTR1 and AGTR2, respectively) and Ang-(1–7) MAS1 proto-oncogene, G protein-coupled receptor (Mas1). Angiotensinogen is the precursor of RAS peptides and is produced by the liver. In the blood stream it is converted into angiotensin I by renin which is released from the juxtaglomerulal cells of kidney due to the drop of blood pressure. Primarily in the pulmonary circulation angiotensin I is further processed by angiotensin-converting enzyme (ACE) into Ang II which binds to AGTR1 and AGTR2 receptors. Elevated Ang II production and AGTR1 activation contributes to elevated blood pressure, aldosterone secretion, water retention, proliferation, and pro-inflammatory and pro-fibrotic responses. AGTR2 receptor activation has the opposite effect, resulting in vasodilation. Ang II is further processed by angiotensin-converting enzyme 2 (ACE2) to a heptapeptide Ang-(1–7) which is an endogenous ligand for vasodilating Mas1 and AGTR2 receptors^4,5^. Besides vasodilatation, the activation of those two receptors results in nitric oxide release, subsequent hyperpolarization via activation of potassium channels, anti-inflammatory and anti-fibrotic responses, inhibits proliferation and finally leads to apoptosis^4^. Deregulation of the RAS system is implicated in the pathology of many diseases, including pulmonary fibrosis^4^, cancer^6^, diabetes^7^, aging and neurodegeneration^8^.

The aim of this study was to evaluate the possible effects of pharmacological treatment with valproate and liraglutide (or their combination) on the regulation of gene expression levels of the key elements of RAS system. Therefore, the expression levels of *Ace, Ace2, Agtr1a, Agtr1b, Agtr2* and *Mas1* were evaluated in the lung tissue of rats treated for eight days with two drugs shown to modify RAS system function: valproate, liraglutide, or their combination.

## Results

To investigate how liraglutide, valproate or their co-treatment can modulate the function of the RAS system, we performed gene expression analyses of the lower lung lobe, extracted from rats after 8 days of treatment. The data were analysed using non-parametric Kruskal-Wallis test followed by Dunn’s multiple comparison test.

Gene expression analyses revealed no statistically significant changes in *Ace* (Figure 1a) and *Ace2* (Figure 1b) expression levels between treatment groups (p > 0.05), although all treatments tended to increase *Ace* mRNA expression compared to saline group and valproate treatment tended to elevate *Ace2* expression. It has previously been shown that *Ace2* and *Ace* are co-expressed in the lung and *Ace* mRNA expression levels are higher compared to *Ace2*^*9*^. Similarly, our results indicate that *Ace* mRNA expression levels are much higher than *Ace2* mRNA level (Figure 1a, b).

**Figure 1.**
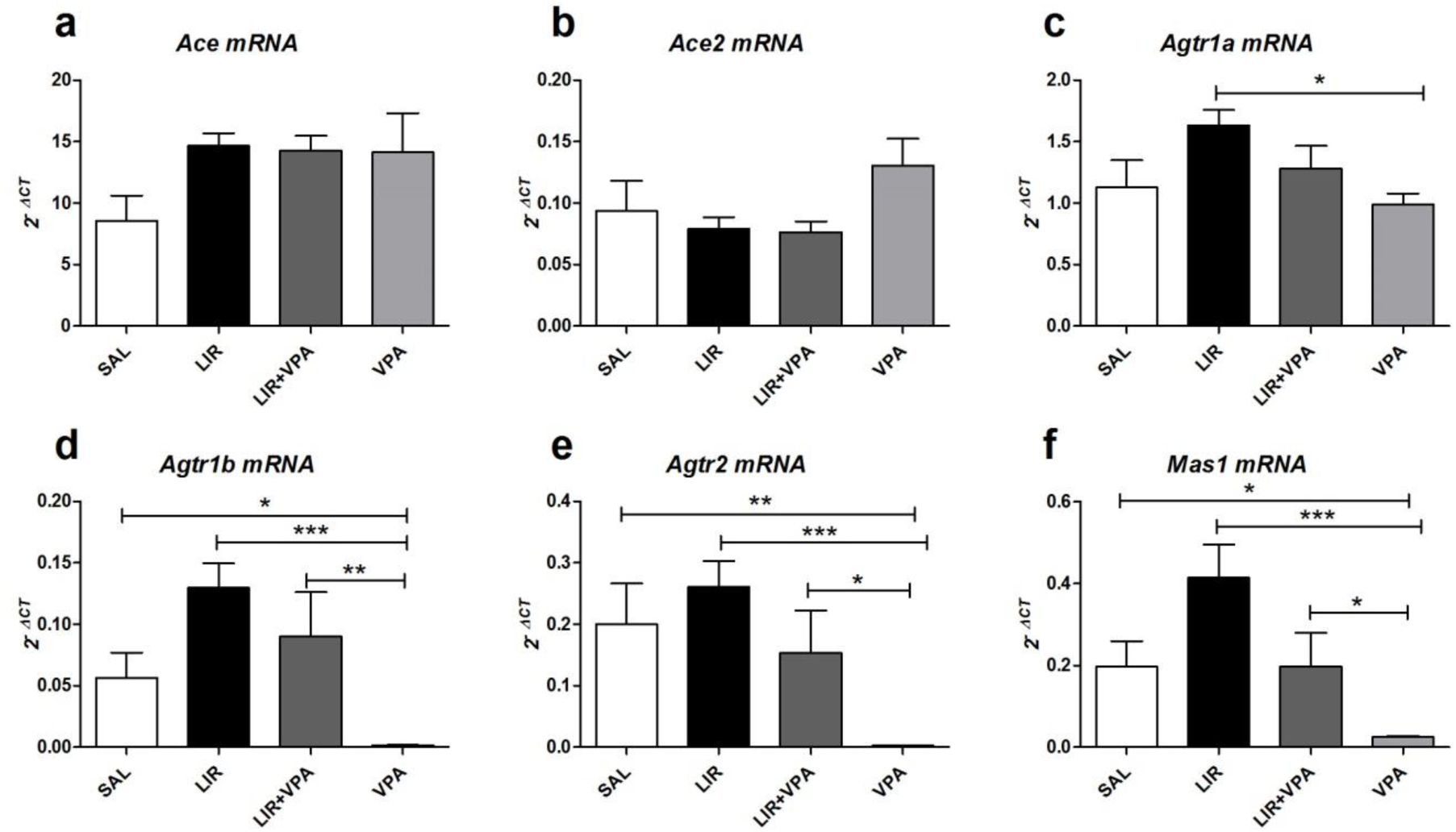
Lung gene expression analyses after 8 days of liraglutide (LIR), valproate (VPA) or liraglutide and valproate (LIR+VPA) co-treatment compared to saline group (SAL). Gene expression of (a) *Ace*; (b) *Ace*2; (c) *Agtr1a*; (d) *Agtr1b*; (e) *Agtr2* and (f) *Mas1*. The data were compared using nonparametric Kruskal-Wallis test followed by Dunn’s multiple comparison test; *p < 0.05, **p < 0.01, ***p < 0.001 compared to saline group. The data are presented as the mean ± SEM, n = 8 per group.

In rodents two Agtr1a receptor subtypes are present, the Agtr1a and Agtr1b encoded by *Agtr1a* and *Agtr1b*, respectively, and they appear to have differential regulation of their expression and possibly different functional properties^10,11^. Therefore, we performed gene expression analyses of both subtypes of the Agtr1 receptors.

Liraglutide had an increasing effect on *Agrt1a* (Figure 1c) mRNA levels compared to saline group, but this difference was not statistically significant (p > 0.05), however, *Agtr1a* gene expression was significantly downregulated after valproate treatment compared with liraglutide treatment (p < 0.05).

*Agtr1b* (Figure 1d) expression was decreased in valproate treatment group compared to saline treated animals (p < 0.05). Liraglutide had an increasing effect on *Agrt1b* mRNA levels compared to saline group, but this difference was not statistically significant (p > 0.05). Additionally, valproate treatment had a significant decreasing effect on *Agrt1b* expression compared to liraglutide treatment (p < 0.001), although valproate and liraglutide co-treatment reversed this decline (p < 0.01).

*Agtr2* (Figure 1e) expression was decreased in valproate treatment group compared to saline treated animals (p < 0.01) similarly to *Agtr1b* expression. Liraglutide had a slight increasing effect on *Agtr2* mRNA levels compared to saline group, even though the difference was not statistically significant (p > 0.05). Most importantly, valproate treatment had a significant decreasing effect on *Agtr2* expression compared to liraglutide treatment (p < 0.001), although in co-treatment liraglutide reversed the suppressing effect of valproate (p < 0.05).

*Mas1* (Figure 1f) expression was decreased in valproate treatment group compared to saline treated animals (p < 0.05) in the same manner as in *Agtr1b* and *Agtr2* expression. Liraglutide had an increasing effect on *Mas1* mRNA levels compared to saline group, but the difference was not statistically significant (p > 0.05). Additionally, valproate treatment had a significant decreasing effect on *Mas1* expression compared to liraglutide treatment (p < 0.001), although valproate and liraglutide co-treatment reversed this decline (p < 0.05).

In conclusion, valproate treatment downregulated the expression levels of *Agrt1b, Agtr2* and *Mas1* basically to zero compared with saline treated animals and interestingly there is a clear pharmacological antagonism of this action by liraglutide (Figure 1d - f).

## Discussion

In the lungs it has been shown that ACE is much more expressed compared to ACE2^2^. Similarly, our results indicate that *Ace* mRNA expression levels are significantly higher than *Ace2* mRNA level (Figure 1e, b).

Liraglutide is a glucagon-like peptide-1 receptor agonist (GLP1 receptor agonist), a common medication used to treat diabetes mellitus type 2 and obesity^12^. GLP1 receptor agonists have been shown to regulate several cellular targets to reduce cellular stress. Importantly, GLP1 receptor agonist treatment was found to upregulate ACE2 expression^1^.

Valproate is primarily used to treat epilepsy and bipolar disorder and to prevent migraine headaches. The precise mechanism of action of valproate is unknown; however, its effects include alteration of GABA levels, blockage of voltage-gated sodium channels, and inhibition of histone deacetylases^13^. Recent results from two *in silico* studies suggest that valproate can downregulate *Ace2* and *Agtr2* gene expression^2,3^. Thus, valproate may modulate the function of RAS system.

Based on this knowledge, we investigated how liraglutide and valproate treatment affect gene expressions of key genes of the RAS system in rat lungs.

Eight-day treatment with liraglutide, valproate or their combination had no major effect on gene expression levels of *Ace, Ace2* or *Agtr1a* mRNA in rat lungs (Figure 1a–c). However, valproate treatment resulted in robust downregulation of *Agtr1b, Agtr2* and *Mas1* mRNA levels in rat lungs (Figure 1d–f): such effect of valproate was clearly pharmacologically prevented by liraglutide (see combination treatment group, Figure 1d–f). Furthermore, liraglutide treatment tended to increase the expression of these genes, however, these effects were statistically non-significant. These results are in agreement with the findings of *in silico* studies where the effect of valproate on modulation of Agtr2 was predicted^2^.

In summary, our results indicate that valproate treatment down-regulates the gene expression of *Agrt2, Agtr1b* and *Mas1* in the rat lungs, and therefore valproate can play an important role in the modulation of the RAS system.

## Materials and Methods

### Animals

For this study, 3.5–4-month-old outbred CD® (Sprague Dawley) rats were used (in house breeding, stock line originated from Charles River Laboratories). Outbred rat line was chosen for better translational value. The animals were housed in groups of four animals per cage under a 12 h light/dark cycle (lights on at 7 a.m.). Rats had unlimited access to food and water. Sniff universal mouse and rat maintenance diet (Sniff #V1534) and reverse osmosis-purified water were used. Experiments were performed between 9 a.m. and 5 p.m.

All experimental protocols were approved by the Estonian Project Authorisation Committee for Animal Experiments (No 165, April 3, 2020), and all experiments were performed in accordance with the European Communities Directive of September 2010 (2010/63/EU).

### Repeated liraglutide and valproate treatment

The rats were randomly allocated to the liraglutide (LIR, n=8), valproate (VPA, n=8), liraglutide + valproate (LIR + VPA, n=8), or saline control group (SAL, n=8). The liraglutide dose was 0.4 mg/kg/day (Novo Nordisk, Denmark) and the valproate dose was 0.3 g/kg/day (valproic acid sodium salt, Sigma-Aldrich P4543). Effective and safe doses of the drugs were chosen based on our previous studies^22, 23^.

All drugs where dissolved in 0.9% saline and injected subcutaneously at the volume of 1 ml/kg. The control group received 0.9% saline (vehicle) subcutaneously every day at the volume of 1 ml/kg. After eight days of treatment and within 4 h after the last injection the animals were killed with intraperitoneal injection of Euthasol vet (dose 300 mg/kg). Lower lobes of the lungs were dissected, washed quickly with 0.9% saline and snap frozen in liquid nitrogen. Tissue samples were stored at -80°C until analysis.

### RNA isolation, cDNA synthesis and gene expression analyses

The lower lung lobe was homogenized using Precellys lysing Kit CK14 (Bertin Instruments) and Precellys homogenizer (Bertin Instruments). RNA from rat lungs was isolated from the lysate using Direct-zol RNA MiniPrep (Zymo Research), according to the manufacturer’s protocol. First-strand cDNA was synthesized using random hexamers and SuperScript™ III Reverse Transcriptase (Invitrogen, USA). qPCR using TaqMan Gene Expression Assays, and Taqman Gene Expression Mastermix (Thermo Fisher Scientific) was used to analyze *Ace* (Rn00561094_m1), *Ace2* (Rn01416293_m1), *Agtr1a* (Rn02758772_s1), *Mas1* (Rn00562673_s1), *Agtr1b* (Rn02132799_s1) and *Agtr2* (Rn00560677_s1) expression. Relative quantification was performed using the 2-ΔCt method, with *Hprt1* (hypoxanthine-guanine phosphoribosyltransferase; Rn01527840_m1) as an internal control.

### Data analysis

The data were analyzed using GraphPad Prism version 5 software (GraphPad Software Inc., San Diego, CA, USA). p < 0.05 was considered statistically significant. The data are presented as the mean ± SEM and were compared using nonparametric Kruskal-Wallis test followed by Dunn’s multiple comparison test.

## Acknowledgements

This research was supported by grant PSG471 (Mario Plaas) from the Estonian Research Council.

## Authors’ contributions

M.P. conceived the study. M.P., E.V., K.S. and A.T. designed the experiments, M.P. directed the study. A.T. and T.J. performed animal experiments. K.S. performed gene expression analysis. K.S., M.P., A.T., and J.I. participated in analysis and interpretation of the data and writing of the manuscript. A.T. and M.P. wrote most of the manuscript. All the authors have read and approved the final version of the manuscript.

